# Major data analysis errors invalidate cancer microbiome findings

**DOI:** 10.1101/2023.07.28.550993

**Authors:** Abraham Gihawi, Yuchen Ge, Jennifer Lu, Daniela Puiu, Amanda Xu, Colin S. Cooper, Daniel S. Brewer, Mihaela Pertea, Steven L. Salzberg

## Abstract

We re-analyzed the data from a recent large-scale study that reported strong correlations between microbial organisms and 33 different cancer types, and that created machine learning predictors with near-perfect accuracy at distinguishing among cancers. We found at least two fundamental flaws in the reported data and in the methods: (1) errors in the genome database and the associated computational methods led to millions of false positive findings of bacterial reads across all samples, largely because most of the sequences identified as bacteria were instead human; and (2) errors in transformation of the raw data created an artificial signature, even for microbes with no reads detected, tagging each tumor type with a distinct signal that the machine learning programs then used to create an apparently accurate classifier. Each of these problems invalidates the results, leading to the conclusion that the microbiome-based classifiers for identifying cancer presented in the study are entirely wrong. These flaws have subsequently affected more than a dozen additional published studies that used the same data and whose results are likely invalid as well.

## Introduction

Bacteria and viruses have been implicated as the cause of multiple types of cancer, including human papillomavirus for cervical cancer (1), *Helicobacter pylori* for stomach cancer (2), and *Fusobacterium nucleatum* for colon cancer (3), among others. However, until a few years ago, little evidence indicated that a complex microbiome–a mixture of various bacteria and viruses– might affect the etiology of other cancer types. This changed after a large-scale analysis of 17,625 samples from the Cancer Genome Atlas (TCGA) reported that, in the sequence data from 33 types of cancer, a distinctive microbial signature was present in 32 of the cancer types (4). These signatures were remarkably accurate at discriminating between each tumor type and all other cancers. For 15 cancer types, signatures were created that could distinguish between tumor and normal tissue, and for 20 cancer types, signatures were developed to identify tumors based on microbial DNA found in the blood of those patients. The machine learning models created in this study had surprisingly high accuracy, with most models ranging from 95-100% accurate.

However, despite efforts taken by Poore *et al*. to remove contaminating species and to avoid common biases like batch effects, we were concerned because many of the machine learning models reported in the study were based on genera that did not make sense in the context of human disease. The models included species that had never been reported in humans, and that were associated only with extreme environments, ocean-dwelling species, plants, or other non-human environments.

Multiple studies over the past decade have reported that the problem of contamination is not limited to the physical samples themselves: in addition, genome databases are contaminated (in a different sense of the term) with large numbers of mis-labeled sequences. The biggest problem, as reported in one study (5), is that human DNA has contaminated the assembled genomes of thousands of bacteria. An even larger study showed that cross-contamination with the wrong species is ubiquitous, affecting over 2 million entries in the GenBank database (6). These contamination events are predominantly present in draft genomes, where some sequences (“contigs”) that originated from humans or other non-microbial species are mis-labeled with the name of a bacterial, fungal, or other microbial species. Database contamination can, in turn, lead to misclassification of human reads that match a contaminated non-human genome.

This contamination problem is of particular concern when using a metagenomics analysis method to classify reads that are derived from a human sample and that have a relatively small number of microbial reads (7-9), which is precisely the scenario encountered in the cancer microbiome study (4). The original samples were collected from human tumors and normal tissue, where the vast majority of the sequenced reads were human. Poore *et al*. reported (4) that 7.2% of the raw reads were classified as non-human, and we were concerned that a substantial fraction of those reads were, in fact, human. Our results below confirm that this concern was legitimate.

## Results

We re-analyzed all of the raw and normalized taxonomic classification data from the Poore *et al*. study (4), which included read counts that were summarized at the genus level. This included the counts for 1993 genera for each of 17,625 samples. Their raw count matrix was created by processing the data with Kraken, a metagenomics classification method developed originally in one of our labs (10, 11). In addition, we downloaded and re-analyzed 1,255 of the original TCGA samples from three cancer types: bladder urothelial carcinoma (BLCA), head and neck squamous cell carcinoma (HNSC), and breast invasive carcinoma (BRCA) (**Table 1**).

**Table 1.** Number of cancer samples downloaded from TCGA for re-analysis, from bladder urothelial carcinoma (BLCA), head and neck squamous cell carcinoma (HNSC), and breast invasive carcinoma (BRCA). The last column shows the number of reads that did not align to the human genome in the TCGA raw BAM files. WGS: whole-genome shotgun sequencing.

|  |  |  |  |  |  |
| --- | --- | --- | --- | --- | --- |
| Cancer type | Number of samples downloaded |  |  | Total read count | Initially unmapped reads |
|  | WGS | RNA-seq | Total |  |  |
| BLCA | 277 | 406 | 683 | 205,521,556,080 | 5,032,358,291 (2.4%) |
| HNSC | 334 | 0 | 334 | 258,961,253,944 | 3,573,898,240 (1.4%) |
| BRCA | 238 | 0 | 238 | 324,824,097,837 | 1,532,210,153 (0.5%) |

### Filtered “non-human” reads contained millions of human reads

As described in Poore *et al*. (4), their analysis began with reads that did not align to known human reference genomes based on the mapping information in the raw BAM files from TCGA.

Those BAM files were the results of using program such as bwa (12) or Bowtie2 (13, 14) to align the reads against a version of the human reference genome, either GRCh37 (hg19) or GRCh38 depending on the date when the samples were processed. This alignment process is imperfect, and many human reads can fail to align from a typical sample. Thus, if one simply downloads the reads that did not align to the human genome, as Poore *et al*. did, many of the reads retrieved in this way will still be human.

To illustrate, we re-aligned all of the initially unmapped reads from the 1,255 BLCA, HNSC, and BRCA samples shown in the last column of **Table 1**. In the original datasets, the proportion of reads that originally did not map to the human genome was 2.4%, 1.4%, and 2.7% respectively. We re-aligned these unmapped reads to the complete CHM13 human genome (15) using Bowtie2 (14), and identified 981,451,972 (19.5%), 519,222,095 (14.5%), and 785,947,157 (51%) additional reads that matched human. This equates to an average number of human reads per sample of 1.39 million, 1.55 million, and 3.3 million in the BLCA, HNSC, and BRCA datasets respectively.

Thus in each of these datasets, the strategy of relying on the mapping information in the raw BAM files, as Poore *et al*. did, leaves on average 1.4–3.3 million human reads in each sample.

### Bacterial read counts were inflated by many orders of magnitude

The presence of millions of human reads in the samples means that in the primary analysis step in Poore *et al*., where they matched all of these reads to a microbial database, any microbial genome that contained short regions of human sequences could generate large numbers of false positive matches; i.e., reads that were reported to match a bacterial genome when in fact the reads were from human DNA. As mentioned above, thousands of draft genomes do indeed contain small amounts of human DNA sequence that are erroneously labeled as bacterial (5).

The TCGA read data were analyzed with the Kraken program (11), a very fast algorithm that assigns reads to a taxon using exact matches of 31 basepairs (bp) or longer. The Kraken program is highly accurate, but it depends critically on the database of genomes to which it compares each read. Poore *et al*. used a database containing 59,974 microbial genomes, of which 5,503 were viruses and 54,471 were bacteria or archaea, including many draft genomes. Notably, their Kraken database did not include the human genome, nor did it include common vector sequences. This dramatically increased the odds for human DNA sequences present in the TCGA reads to be falsely reported as matching microbial genomes. This problem can be mitigated by including the human genome and by using only complete bacterial genomes in the Kraken database.

### Re-analysis of bladder cancer samples

We re-analyzed 156 bladder cancer samples (all WGS primary tumor and normal tissue samples from BLCA) by matching them against a curated, publicly available Kraken database (16) that contained only finished bacterial genomes as well as viruses, eukaryotic pathogens, the human genome, and commonly-used laboratory vectors (see Methods). None of the bacterial genomes in this database were draft genomes. We first re-filtered the unmapped reads by aligning them to the human CHM13 reference genome, and we only analyzed reads remaining after this second filtering step (see Methods). Note that even with two rounds of alignment against the human genome, many of the reads in each sample were still classified as human by the Kraken program using our database.

Figure 1. and Supplementary Table S1 show the top 20 most-abundant microbial genera as reported in the Poore *et al*. study for BLCA, compared to the read counts found in our analysis. As shown in the figure, the top genera from Poore *et al*. were *Streptococcus, Mycobacterium*, and *Staphylococcus*, with average read counts per sample of 560K, 411K, and 241K respectively. In our re-analysis of the same samples, we found far fewer reads in these genera: an average of 36, 6, and 282 reads respectively, values that are 16,000, 67,000, and 854 times smaller. Table S2 shows the top 20 genera found in our analysis, which had abundances ranging from 12–447 reads per sample.

**Figure 1.**
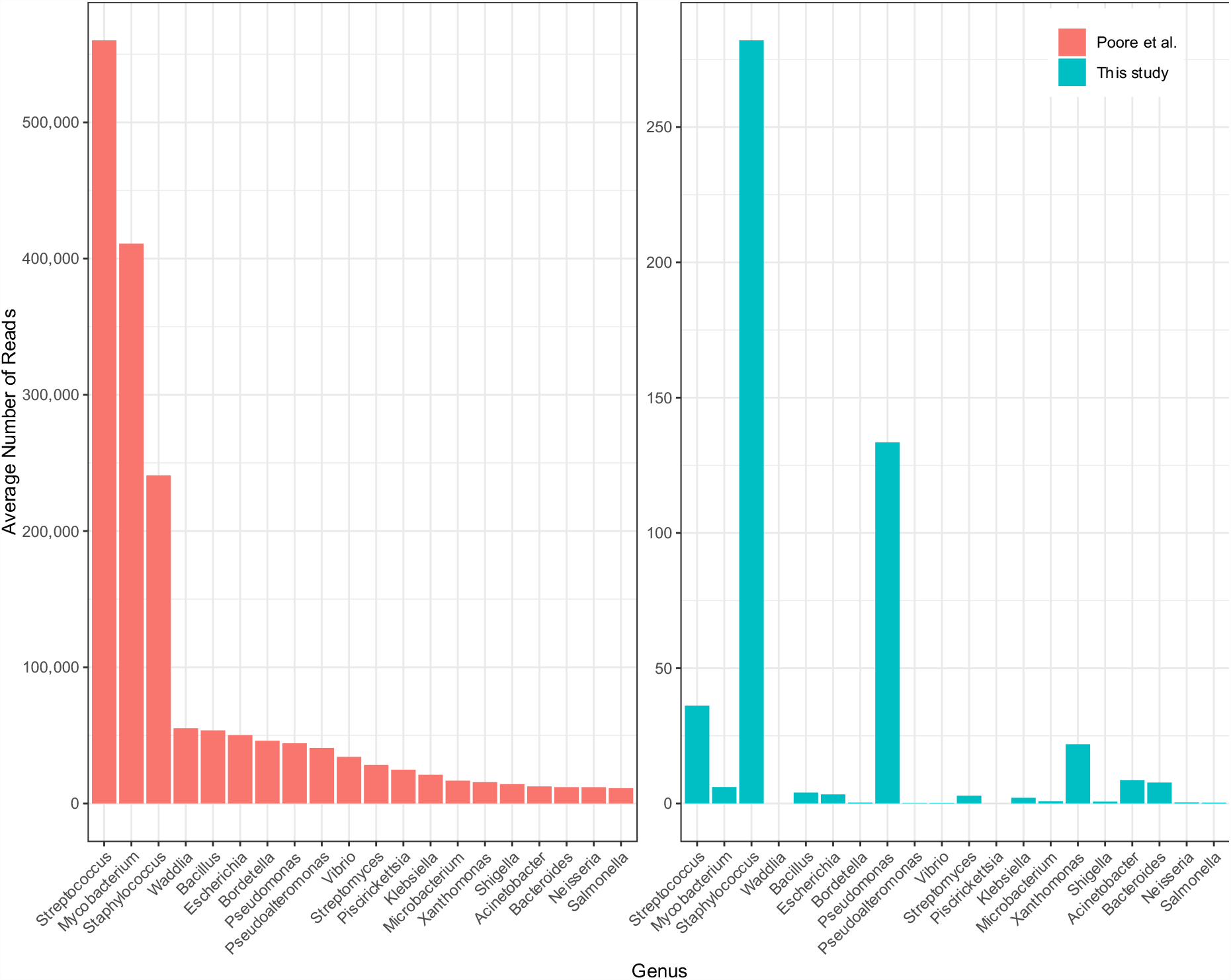
Average number of reads per sample in bladder cancer (BLCA) in the top 20 most-abundant genera reported in Poore *et al*. (left), averaged across 156 whole-genome sequencing samples. On the right are the counts for the same samples and same genera, in the same order, as computed in our re-analysis. Note that the y-axis scales are different by a factor of 2000. The x-axis shows genus names.

As we describe below, the vast majority of the excessive counts in the Poore *et al*. study were apparently due to human reads in the filtered data that were incorrectly labeled as bacterial.

Because filtering the raw reads only against GRCh37 or GRCh38 did not remove all human reads, the input to their metagenomics pipeline included 1.4–3.3 million human reads per sample, and these reads explain the dramatic over-counts shown on the left side of Figure 1.

We also compared the read counts for the genera that were given the highest weights in the machine learning models created by Poore *et al*. Supplementary Tables S3–S4 show the average read counts for the 20 top-weighted genera in models that classified bladder cancer tumors versus other tumor types and tumors versus normal tissue. In our analysis, nearly all counts for these genera averaged between 0 and 1, with a maximum value of just 18 reads (for *Campylobacter*). Nearly half of the top-weighted genera had average read counts below 10 in the Poore *et al*. data as well, although several had counts in the thousands. Below we explain how genera with raw counts near zero were selected by the machine learning models.

### Re-analysis of head and neck cancer and breast cancer samples

We conducted the same re-analysis on 334 HNSC samples and 238 BRCA samples (see Methods). As with the BLCA samples, we filtered to remove reads matching the CHM13 human genome and then used the Kraken program to match all reads against a curated database of microbes, common vectors, and the human genome.

Supplementary **Figures S1-S2** and Tables S5-S6 show the read counts for the 20 most abundant genera in the HNSC data (Fig. S1) and BRCA data (Fig. S2) as computed by Poore *et al*., contrasted with our read counts for the same genera in the same samples. As with the bladder cancer results shown in Figure 1, the average read counts reported by Poole *et al*. in both cancer types were consistently *hundreds or thousands of times higher* than found in our analysis. For most of these putatively abundant genera in all three cancer types, we found average read counts near zero, while Poore *et al*. reported read counts ranging from tens of thousands to over one million. As we demonstrate below, the vast majority of these over-counts are human reads that were erroneously assigned to bacteria.

### Nearly all of the raw numbers in the Poore et al. study are incorrect and far too high

We then looked more broadly at all of the read counts, for all genera, for the BLCA, HNSC, and BRCA whole-genome samples, focusing on the non-zero counts reported by Poore *et al*. Note that in the original raw data matrix, 21,074,259 (60%) of the entries were zero and 14,071,985 (40%) were non-zero. Each cell in the matrix represents a sample/genus pair; i.e., the count of the number of reads from a given sample that were assigned to a given genus.

**Table 2** summarizes our comparisons. For every non-zero genus in every sample, we compared the number of reads reported by Poore *et al*. to the number we found in our analysis. The table focuses on sample/genus pairs where the read counts were at least 10, on the assumption that smaller values likely represent noise or contamination.

**Table 2.** Microbial read totals found by Poore *et al*. (4) for three cancer types, compared to counts computed in a re-analysis using a database with only complete bacterial genomes. BLCA: bladder cancer; HNSC: head and neck cancer; BRCA: breast cancer.

|  |  |  |  |  |
| --- | --- | --- | --- | --- |
| Cancer type | Read counts reported in Poore *et al.* | Sample/genus pairs | Poore *et al.* value is |  |
| | | | $\geq 10$ times too high | within 50% of re-estimate |
| BLCA samples (156 total) | $\geq 10$ reads | 37,723 | 37,174 (98.5%) | 90 (0.02%) |
| | $\geq 100$ reads | 17,028 | 16,928 (99.4%) | |
| HNSC samples (334 total) | $\geq 10$ reads | 114,640 | 106,544 (92.9%) | 2,152 (1.88%) |
| | $\geq 100$ reads | 55,259 | 52,689 (95.3%) | |
| BRCA samples (238 total) | $\geq 10$ reads | 83,476 | 81,736 (97.9%) | 334 (0.4%) |
| | $\geq 100$ reads | 46,025 | 45,686 (99.3%) | |

As shown in the table, in the BLCA samples the number of reads reported by Poore *et al*. was at least *10 times larger* than our results for 98.5% of the data entries. If we looked only at samples and genera where Poore *et al*. found ≥ 100 reads, their value was more than 10 times too large in 99.4% of all cases. The results were similar for HNSC, where 97.9% of values were at least 10 times too high, and in BRCA where 97.9% of the read counts were inflated at least 10-fold.

We also computed how many of the non-zero read counts were at least approximately the same as the value as determined in our re-analysis. In the BLCA samples, only 90 out of 37,723 (0.02%) were within 50% of the counts that we found in the same samples. Equivalently, fewer than 1 in 400 non-zero values in the bladder cancer data were within 50% of the value found upon re-analysis. The HNSC and BRCA read counts were only marginally better, with just 1.9% and 0.4% respectively within 50% of the correct value. Thus the vast majority of the non-zero data in the original data matrix of Poore *et al*.–the data upon which all of their results were based–appears to be wrong, by very large amounts.

### How human reads create the false appearance of bacteria

The likely reason for these vast over-counts is that human reads were erroneously categorized as bacterial by Poore *et al*. The number of human reads matching bacteria was unrelated to the actual presence of bacteria in the tumor sample; instead, it was determined by the database itself, in which many draft bacterial genomes contained mis-labeled human sequences.

To illustrate how such high read counts can appear when few or even no reads from a bacterial genus are present, we did a deep analysis of two genera, *Streptococcus* and *Waddlia*, in one primary tumor sample, s2707 (case ID TCGA-DK-A1AB), from the BLCA data. We chose these genera because they were reported to be among the most abundant in Poore *et al*., as shown in Figures 1, S1, and S2. Sample s2707 was reported by Poore *et al*. to have 327,985 *Streptococcus* reads and 20,673 *Waddlia* reads. When we aligned s2707 to our Kraken database, which only contains complete bacterial genomes, we found just 1 read labeled as *Streptococcus* and none labeled as *Waddlia*.

We then extracted all reads from s2707 that did not match the human genome in the TCGA BAM file, which comprised 11,997,726 unmapped reads. Next we built a custom Kraken database containing all 10,270 *Streptococcus* genomes available in GenBank as of 2016. (We chose 2016 because Poore *et al*. downloaded all bacteria for their database in June of 2016 (4).) We built a second Kraken database that contained all four of the *Waddlia* genomes that are publicly available. We then ran KrakenUniq (11) to map the ∼12 million unaligned reads from s2707 against both databases, and found that 1,434,287 read pairs were classified as *Streptococcus* and 197,811 as *Waddlia*, respectively. This finding demonstrates that it is indeed possible, starting with the unaligned reads from a cancer sample (s2707), to find large numbers of reads from each of these genera when aligning against a database built entirely from bacterial genomes, as long as that database does not contain the human genome.

To confirm that the over-counts were due to human reads that erroneously matched bacteria, we then extracted all reads labeled as either *Streptococcus* or *Waddlia* in the Kraken analyses above and aligned them to the CHM13 human genome using Bowtie2 (14). This step revealed that 98.1% and 98.9% of the reads (respectively) matched human DNA. Thus the Kraken matches were nearly all false positives, caused by the presence in the database of bacterial genomes that (erroneously) contained human sequences.

Finally, to emphasize the effect of omitting the human genome from the Kraken database, as Poore *et al*. did, we created two more databases: one containing the 10,270 *Streptococcus* genomes plus human, and one with the 4 *Waddlia* genomes plus human, using the CHM13 human genome in both cases. We then classified all reads from sample s2707 again. When classified against the first database, the number of Streptococcus reads dropped from 1,434,287 to 10,792, a 132-fold decline. When using the second database, the number of Waddla reads dropped from 197,811 to 174, a decline of more than 1000-fold.

### Normalization of the reads erroneously created a distinct signature of each cancer

The second major error in the Poore *et al*. study occurred during normalization of the raw read counts. Poore *et al*. used normalized rather than raw data to build all of their machine learning classifiers, in order to remove batch effects (4). In the process of converting the raw counts to normalized values, many of the cancer types (e.g., all tumor samples for one cancer type, or all healthy samples for another cancer type, etc.) were erroneously tagged with distinct values, marking the cancer samples even when the raw values were not informative. The machine learning programs were then able to use these artificial tags to create near-perfect classifiers. We examined the top genera used in many of these classifiers and found numerous examples of this erroneous marking, a few of which are shown here.

First, consider the values for *Hepandensovirus* in Adrenocortical carcinoma (ACC). All of the ACC cancer samples had raw read counts of zero for this virus, but during normalization, 71 of the 79 samples (90%) were assigned the value 3.078874655 by Poore *et al*. Out of all 17,625 samples across all cancer types (including 13,883 primary tumor samples), only 77 other samples had a value equal to or smaller than this value in the normalized data. In the raw data, however, 17,624 samples had zero *Hepandensovirus* reads, and one sample had 2 reads.

As illustrated in **Figure 2**, the extremely non-random distribution of normalized values–all but one of which started as raw values of zero–makes it easy for a machine learning classifier to separate the ACC samples from other cancers. If we call the normalized *Hepandensovirus* value H_N_, then if the model splits the samples using the simple rule H_N_>3.078874655, it will label 71/79 (90%) of the positive samples correctly, and only make 77/17,625 (0.4%) errors (**Figure 2**). This explains why *Hepandensovirus* was the highest-weighted feature for the machine learning model distinguishing ACC from other cancers in the most stringent decontamination (MSD) data set, despite the fact that only 1/17,624 samples had any reads at all matching this virus.

**Figure 2.**
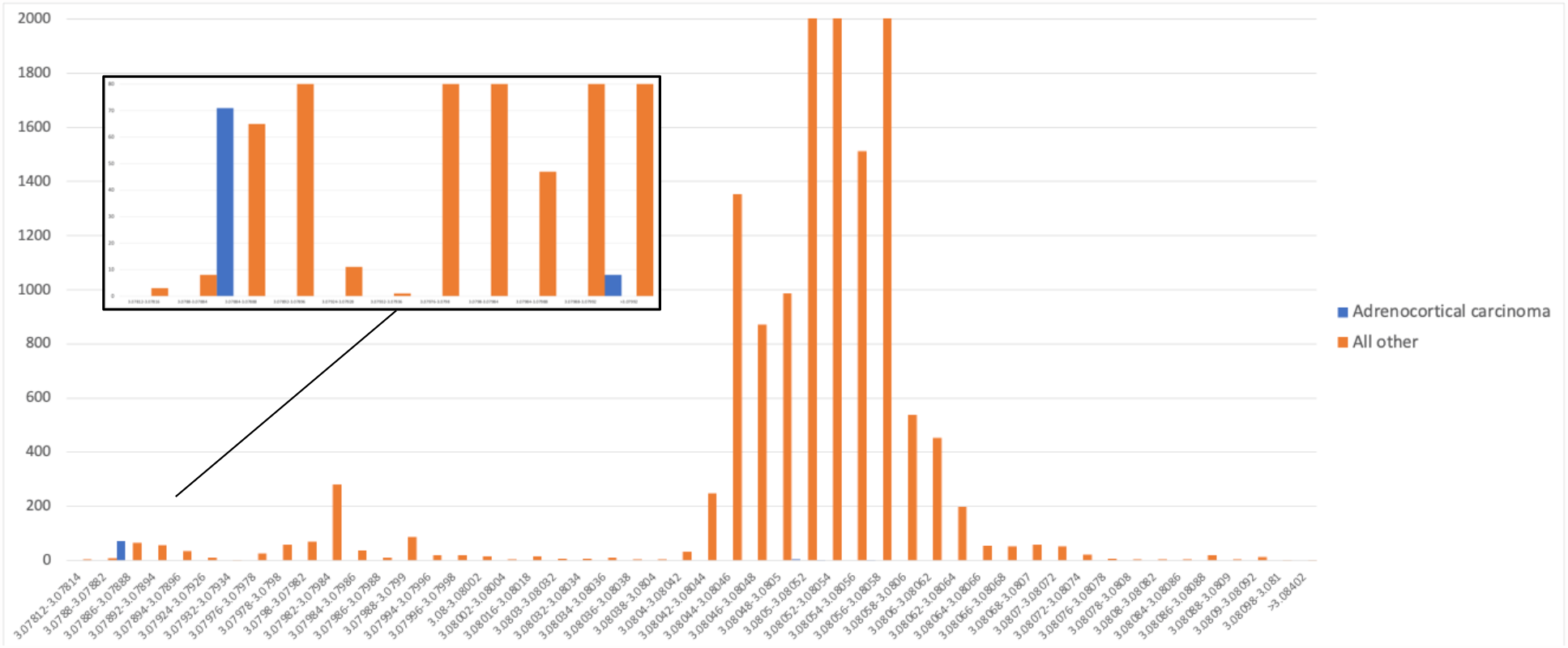
Distribution of normalized counts for *Hepandensovirus* for Adrenocortical carcinoma (blue) versus all other samples (orange). Inset shows zoomed-in view of the distribution for the smallest values. All raw values were zero.

We observed a similar pattern in the normalized values of another genus, *Thiorhodospira*, for the Kidney chromophobe (KICH) tumor samples (**Figure 3**). *Thiorhodospira* was the highest-weighted feature for the machine learning classifier that distinguished KICH from normal tissue in several different models (including the Full dataset, the “likely contaminants removed” (LCR) dataset, and the “all putative contaminants removed” (APCR) dataset). The TCGA-KICH data contained 51 tumor samples and 41 normal tissue samples, and in the raw data, 85 samples had read counts of zero, and 7 samples (4 cancer, 3 normal) had counts of 1 for *Thiorhodospira*, meaning that it had almost no utility as a discriminating feature. In the normalized data, though, the cancer samples were assigned an almost perfectly disjoint set of values from the normal tissues, as shown in Figure 3. Thus once again, the normalization process created an artificial signal separating tumor from normal tissue.

**Figure 3.**
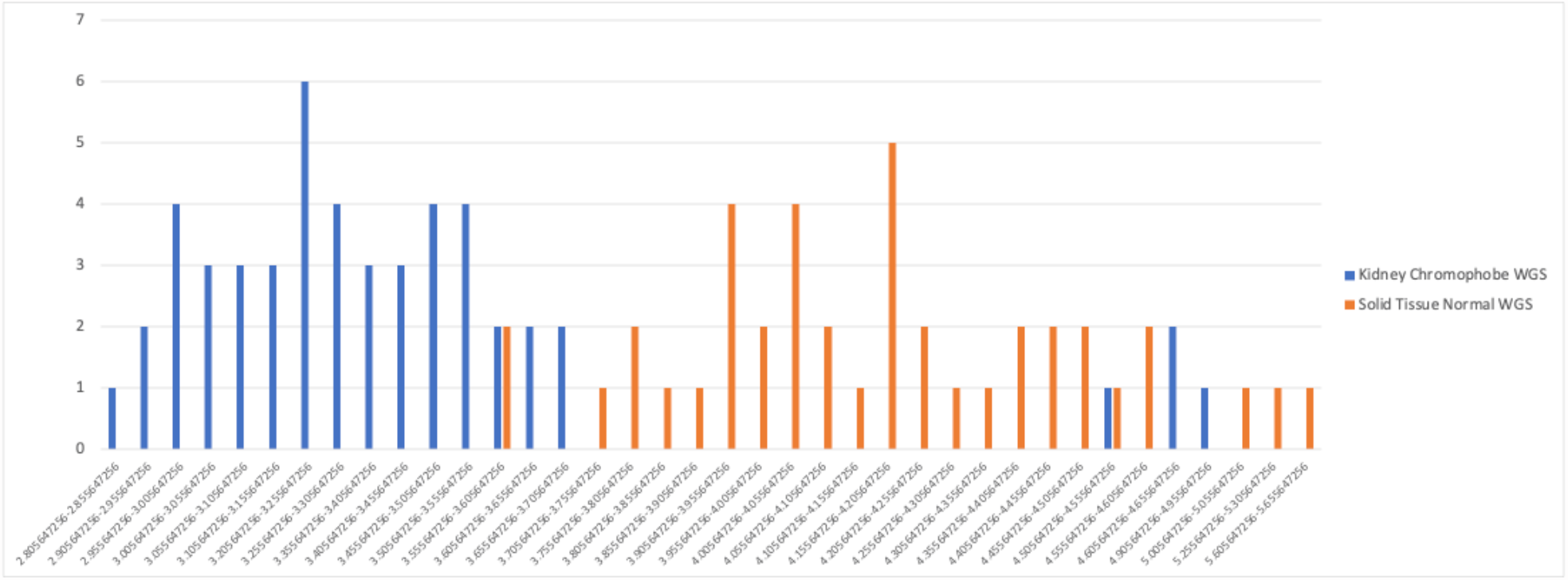
Distribution of normalized counts for *Thiorhodospira* reads in kidney chromophobe (KICH) cancer (blue) and normal (orange) samples. Nearly all raw values were zero except for 7 samples with a raw count of 1.

Another example is *Nitrospira*, which was a highly-weighted genus for the machine learning models in 13 different cancer types in Poore *et al*., including lung squamous cell cancer (LUSC) where it was the top-ranked genus. **Figure 4** shows the normalized counts of *Nitrospira* reads, after Voom-SNM normalization, in the LUSC samples compared to all other cancer types. In the figure, the frequencies of *Nitrospira* in LUSC are shifted to the right; i.e., they have larger values on average than other cancers. This explains why the machine learning model gave *Nitrospira* the highest weight; however, in the raw data there is no such shift to the right. Thus the Voom-SNM normalization process created a signature of lung cancer even though no such signature was present in the original read counts.

**Figure 4.**
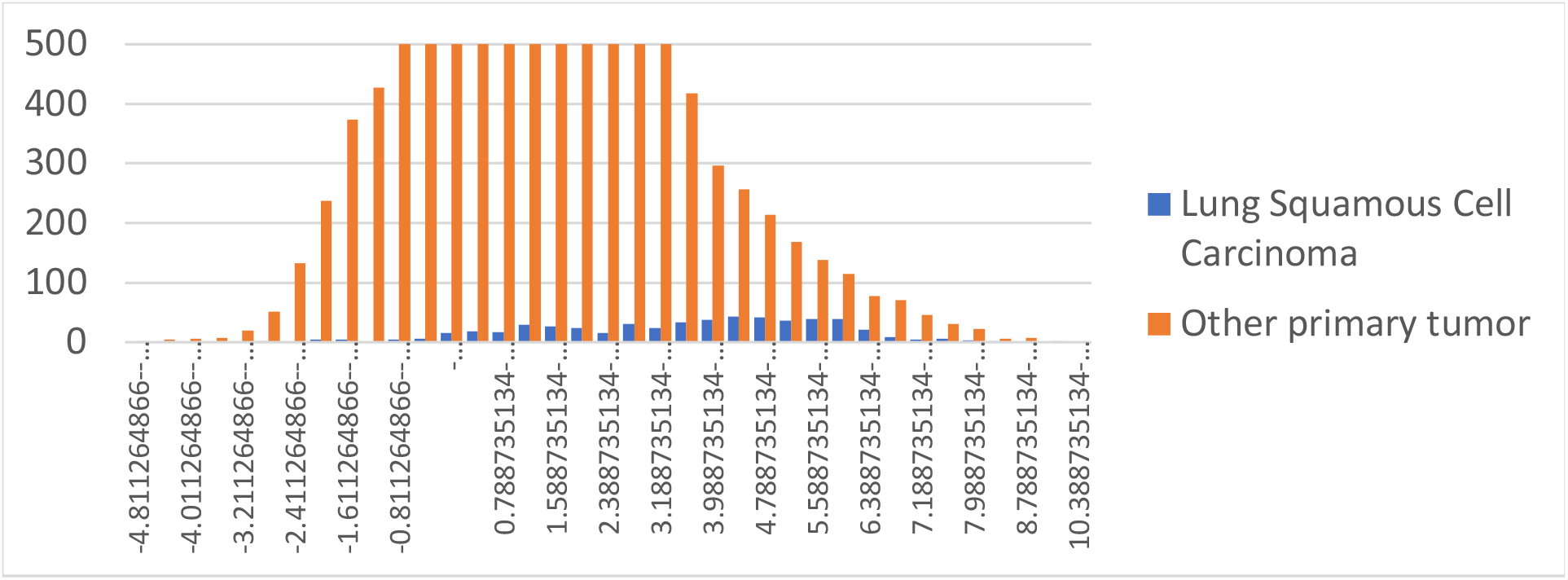
Distribution of normalized read counts in the APCR data set for *Nitrospira* reads found in lung squamous cell carcinoma (blue) and all other cancer types (orange). For clarity, the y-axis is truncated at 500, but the peak of the distribution for other cancers (organe) is at 1389.

We observed this phenomenon again in HNSC, where the genus with the highest weight in the MSD dataset was *Mulikevirus*. This genus had the highest weight both for distinguishing tumor from normal tissue, and for distinguishing HNSC from all other cancers. *All* 906 HNSC samples, including tumor, blood, and normal tissue, had zero reads in the raw data for *Mulikevirus*, making this virus useless at discriminating between tumor and normal samples.

However, in the Voom-SNM normalized data, values for the 70 normal samples were set to lower values than any of the tumor samples, as shown in **Figure 5**. In particular, 38 samples had the identical normalized value of 3.07584214, 18 others had the value 3.07585718, and 5 had the value 3.076237397. The vast majority of the 693 tumor samples had larger values, as shown in the figure. Thus a machine learning model using this genus alone would have very high accuracy, which explains the very high weight given by the model to *Mulikevirus*, despite the fact that all the raw read counts were zero.

**Figure 5.**
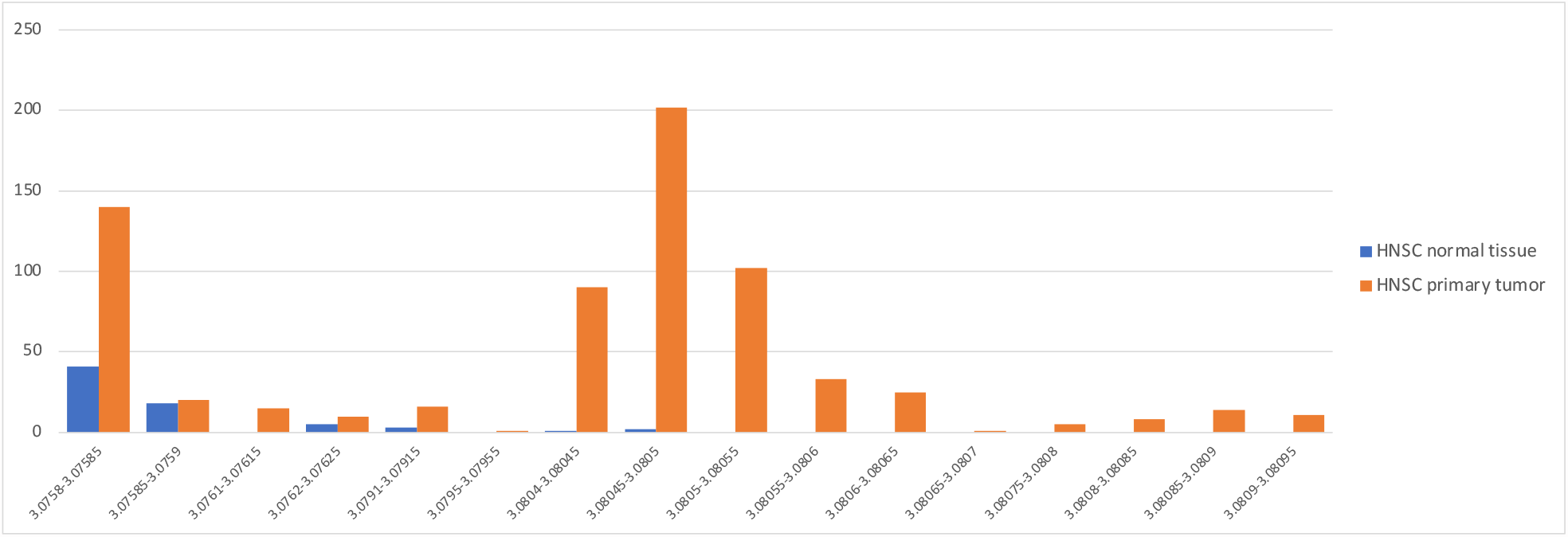
Distribution of normalized counts for *Mulikevirus* reads in head and neck squamous cell (HNSC) cancer (orange) and normal (blue) samples. All raw values were zero.

### Replicating highly accurate classifiers on information-free raw data demonstrates flaws in the normalization process

Given that individual genera such as *Hepandensovirus* were erroneously tagged with tumor-type specific values, we wanted to explore how this tagging would affect the performance of machine learning classifiers on a larger selection of tumor types and taxa. To investigate this question, we extracted a completely empty microbial-sample matrix (all zeros) from the raw Kraken classification data provided in the Poore *et al*. study. To obtain the empty matrix, we filtered the data to retain only genera present in fewer than 50 samples, and then removed any samples with non-zero values for any genus. This produced a matrix containing 16,567 samples and 170 genera in which all values were zero. No machine learning classifier can use such data to discriminate among cancer types, because every entry in the matrix is identical.

We then populated each cell in the empty matrix with its corresponding value from the Voom-SNM normalized data. For this experiment, we used the Voom-SNM data from the “most stringent decontamination” (MSD) dataset, which included only 66 of the 170 taxa in our initial empty matrix. We then filtered to retain only primary tumor samples (*N*=12,803) so that we could attempt to build classifiers discriminating each cancer type from all others.

We then applied the original code provided by Poore *et al*. (4) to classify one tumor type versus all others, and created classifiers for all 32 cancer types using this 12,803×66 matrix. Accuracies for these classifiers are shown in **Figure 6**. Nearly all the models obtained very high accuracy, including a median (across all cancer types) sensitivity of 0.94, a median specificity of 0.9, and a median negative predictive value of 1.0. Several models obtained high positive predictive values (PPV) as well, including those for Stomach Adenocarcinoma (PPV=0.65), Ovarian Serous Cystadenocarcinoma (PPV=0.91), and Glioblastoma Multiforme (PPV=0.92). Comparing model performance between these models and those reported in Poore *et al*., 14 out of 32 models had equal or improved accuracy as measured by area under the sensitivy-specificity curve.

**Figure 6.**
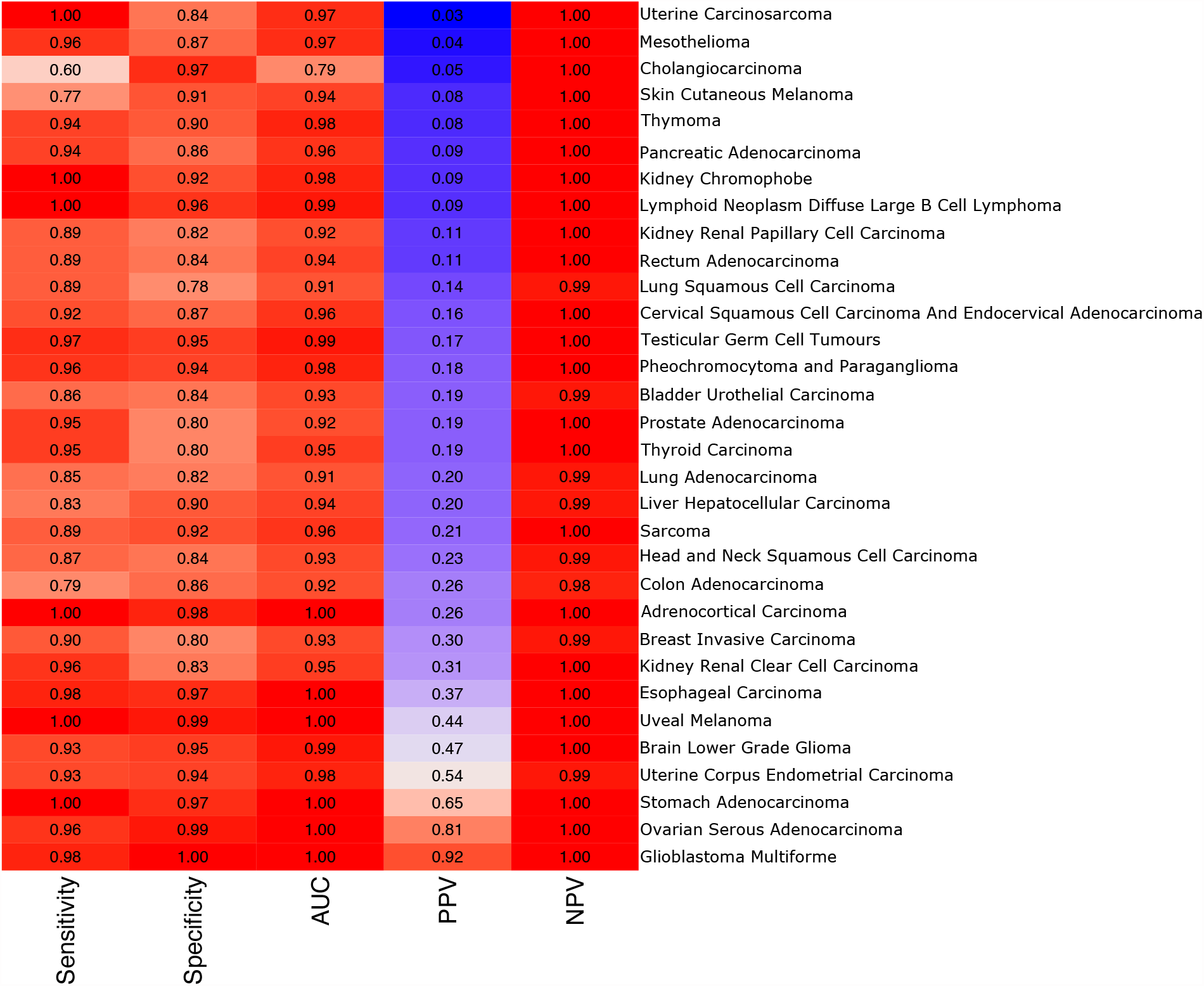
Accuracies for one-vs-all tumor classification models obtained from a selection of samples and genera with zero classified reads prior to normalization. Each row shows the accuracies of a classifier that distinguished one cancer type from all other cancer types in the table. AUC: maximum measured area under the sensitivity-specificity curve. PPV: positive predictive value. NPV: negative predictive value.

Thus despite the fact that the original raw data contained values of zero for all genera and all samples, remarkably high classification accuracy was obtained by the machine learning classifiers, similar to the performance reported in Poore *et al*. All of the signal in these recreated models therefore must be artifactual, arising purely from the Voom-SNM normalization process, which the machine learning methods exploited to create highly accurate classifiers despite the absence of any true signal.

We conclude that the Voom-SNM normalization, at least in the manner employed by Poore *et al*., inadvertently attached prior information about the tumor type to the normalized data. Note that we do not know precisely where Poore *et al*. went wrong in applying the normalization code, but because we have the original read count data and the resulting normalized data, we know that the transformation created the artificial signals that we describe here.

This result not only casts doubt on the claim that tumor types can be distinguished based on a microbial signature, but it also raises concerns about the machine learning models that distinguished between tumor and normal tissue, and those based on microbial reads detected in blood samples.

### Multiple other studies rely on the same flawed data

Since the publication of the study by Poore *et al*. study, more than a dozen studies have downloaded and used the Poore *et al*. data to find additional associations with the cancer microbiome, associations that in each case are likely to be invalid, because the underlying data is invalid. These include the following.

Hermida *et al*. (17) built predictive models for cancer prognosis in multiple cancer types using the Voom-SNM normalized data from Poore *et al*., which they used as the basis for creating machine learning models to predict overall survival and progression-free survival for different cancer types. As shown above, the Voom-SNM data was flawed, introducing a distinctive signature into each cancer type even when the original read counts were all zeros. Thus no classifiers based on this data can be considered valid.

A 2023 study by Parida *et al*. (18) reported finding distinct microbial communities in breast tumors from Asian, black, and white women, based on the raw data matrix downloaded from Poore *et al*., which as described above has vastly over-inflated counts for nearly all genera. A number of the taxa highlighted as being important in this study are extremophiles (e.g., *Halonatronum* and *Salinarchaeum*) that are unlikely to be present in human samples.

Mao *et al*. (19) used the Voom-SNM normalized microbiome data from Poore *et al*. to create a predictive model of survival in breast cancer, based on the abundances of 94 genera. The study claims that its 15-microbe signature can predict overall survival and progression-free survival, but the model includes genera that are not known to exist in humans. For example, one genus is *Methanothermus*, an extremophile archaeon that lives in deep-sea hydrothermal vents at very high temperatures. This genus is extremely unlikely to be present in human breast cancers, and indeed no reads from this genus were found in our re-analysis.

Multiple other studies, including Luo *et al*. (20), Zhu *et al*. (21), F. Chen *et al*. (22), Narunsky-Haziza *et al*. (23), C. Chen *et al*. (24), Lim *et al*. (25), Bentham *et al*. (26), Y. Kim *et al*. (27), Y. Xu *et al*. (28), and Y. Li *et al*. (29), have also utilized the Voom-SNM data from Poore *et al*. to explore various aspects of the tumor microbiome and its potential associations with cancer. However, given the aforementioned flaws and inaccuracies in the Voom-SNM data, caution should be exercised when interpreting the results of any of these studies.

## Discussion

The original findings of a strong association between microbial species and 33 different cancer types were based on a large collection of DNA and RNA sequencing samples taken from human cancers and from matched normal tissues, which in turn was processed by a sophisticated machine learning method to create highly accurate classifiers that could distinguish among tumor types and could distinguish tumor from normal tissue (4). Many of these classifiers used bacterial and viral genera that were not known to exist in humans, and therefore raised questions about their plausibility (30); however this observation alone was not a fatal flaw. It did lead us to explore the machine learning models more closely, though, in an effort to determine why organisms such as non-human extremophile microbes appeared as key features in the classifiers.

After re-analyzing all of the raw and transformed data, and after downloading and re-analyzing the original reads from more than 1,200 tumor and normal samples, we identified two major errors: first, the raw read counts were vastly over-estimated for nearly every bacterial species, often by a factor of 1000 or more. The likely cause of these over-estimates was that the metagenomics database included thousands of draft genomes, which are known to be contaminated with human sequences. Consequently, as we showed above, millions of human reads were erroneously assigned to bacterial or archaeal genera. Second, the process of transforming the raw read counts into normalized values erroneously tagged many of the genera with values that were unique to specific cancer types. When these values were fed to machine learning classifiers, the algorithms discovered these artificial tags and built highly accurate classifiers, often using features (genera) that in the raw data had zero discriminative power. This error seems to have involved every tumor type and many genera that had zero or near-zero read counts across all of the human samples.

Either of these two errors suffices to invalidate the conclusions of the Poore *et al*. study and of the other studies that relied upon the same data. The original data matrix of raw read counts contained millions of wildly inaccurate values, and the normalized data compounded this error by tagging the cancer types with distinctive normalized values. Our conclusion after re-analysis is that the near-perfect association between microbes and cancer types reported in the study is, simply put, a fiction.

## Methods

We downloaded raw reads from the Genome Data Commons at the U.S. National Cancer Institute (gdc.cancer.gov) for three types of cancer from the TCGA project: bladder cancer, head and neck cancer, and breast cancer. For TCGA-BLCA (Bladder Urothelial Carcinoma) we downloaded read data from 683 samples, which included 277 whole-genome shotgun (WGS) samples and 406 RNA-seq samples (**Table 1**). We focused our re-analysis on the WGS samples, which included 129 primary tumor and 27 solid-tissue normal samples. All reads had been previously aligned by the TCGA project to either GRCh38 or GRCh37/hg19 using bwa (12). We extracted all unmapped reads and re-aligned them against the CHM13 human using Bowtie2 (14) to remove additional human reads, and created new files for further downstream analysis.

For TCGA-HNSC (Head and Neck Squamous Cell carcinoma), we downloaded the raw reads from 334 WGS samples, which included 24 solid-tissue normal, 140 blood-derived normal, and 170 primary tumor samples. As with BLCA, we focused the analysis on WGS samples. For TCGA-BRCA (Breast Invasive Carcinoma) we downloaded the unmapped reads from all 238 available WGS samples in TCGA (Table 2), which included 114 primary tumors, 106 blood-derived normal, 16 solid-tissue normal, and 2 metastatic samples. For both HNSC and BRCA, we ran the same two-pass filtering as for the BLCA samples, re-aligning all unmapped reads against CHM13.

Using these two-pass filtered files, for all samples in the BLCA, HNSC, and BRCA data, we ran KrakenUniq (11) against a customized database built from all complete genomes of bacteria and viruses from RefSeq that contained 46,711 bacterial genomes (5981 species), 13,011 viral genomes (9905 species), and 604 archaeal genomes (295 species). It also included a collection of 246 eukaryotic pathogens from EuPathDB (31), the UniVec set of standard laboratory vectors from NCBI (https://www.ncbi.nlm.nih.gov/tools/vecscreen/univec/), and the GRCh38 human genome. This 384GB KrakenUniq database is available for download from https://benlangmead.github.io/aws-indexes/k2. Files containing lists of all species, genera, and NCBI accession numbers in this database are available as Supplementary data files 1–3. All supplemental files and tables from this study are available at https://github.com/yge15/Cancer_Microbiome_Reanalyzed.

Supplemental Tables S8, S9, and S10 contain the read counts at the genus level for all non-zero bacteria, archaea, and viruses found in our re-analysis of the BLCA, HNSC, and BRCA data and summarized in Table 2. Note that even though these numbers are far smaller than those reported in Poore *et al*., they likely still contain some false positives and should be regarded as upper bounds on the actual number of reads from each genus. Supplemental Table S11 contains the top 25 genera identified by the machine learning classifiers created in the Poore *et al*. study, downloaded from http://cancermicrobiome.ucsd.edu/CancerMicrobiome_DataBrowser. These include all the classifiers for the APCR and MSD datasets (separate classifiers were created for each dataset, and for those datasets, the table includes the top genera used for classifying one cancer type versus all others and for distinguishing tumor from normal tissue,

## Acknowledgements

SLS, PG, JL, DP, and MP acknowledge support from the U.S. NIH under grants R01 HG006677 and R35-GM130151. AG, CSC, and DSB acknowledge support from Prostate Cancer UK (MA-ETNA19-003), Big C Cancer Charity (ref 16-09R), The Bob Champion Cancer Trust, and Cancer Research UK.

## Author contributions

AG, YG, JL, DP, AX, and SLS analyzed the data. AG ran the replication studies that created classifiers on the empty data matrix. DP downloaded and re-aligned raw data from TCGA. YG and JL ran metagenomics analyses on the raw cancer data. SLS and AG conceptualized the overall study and wrote the initial manuscript. DSB, CSC, and MP were involved in the conception and design of the study. AG, SLS, CSC, DSB, and MP edited and revised the manuscript and critiqued the output for important intellectual content.

## Conflicts of interest

CSC, DSB and AG are coinventors on a patent application (UK Patent Application No. 2200682.9) from the University of East Anglia/UEA Enterprises Limited regarding the application of biomarker bacterial genera in prostate cancer. All other authors declare no conflicts.

## Notes

https://github.com/yge15/Cancer_Microbiome_Reanalyzed

### Competing Interest Statement

The authors have declared no competing interest.

