## Supplementary materials for "Major data analysis errors invalidate cancer microbiome findings"

### Supplementary Tables and Figures

**Table S1**. Average number of reads in the top 20 most abundant genera for TCGA-BLCA as computed in the Poore *et al*. study, averaged over 129 WGS primary tumor and 27 solid tissue normal samples.

| Genus | Average read count per sample, Poore *et al.* | Average read count per sample, this study |
|---|---|---|
| *Streptococcus* | 560212 | 36 |
| *Mycobacterium* | 410968 | 6.1 |
| *Staphylococcus* | 240880 | 282 |
| *Waddlia* | 55280 | 0 |
| *Bacillus* | 53659 | 4 |
| *Escherichia* | 50207 | 3.4 |
| *Bordetella* | 46075 | 0.4 |
| *Pseudomonas* | 44186 | 133 |
| *Pseudoalteromonas* | 40837 | 0.2 |
| *Vibrio* | 34216 | 0.2 |
| *Streptomyces* | 28317 | 2.9 |
| *Piscirickettsia* | 24818 | 0 |
| *Klebsiella* | 21069 | 2.1 |
| *Microbacterium* | 16773 | 0.9 |
| *Xanthomonas* | 15681 | 22 |
| *Shigella* | 14208 | 0.7 |
| *Acinetobacter* | 12562 | 8.6 |
| *Bacteroides* | 12013 | 7.7 |
| *Neisseria* | 11988 | 0.4 |
| *Salmonella* | 11234 | 0.3 |

**Table S2**. Average number of reads in the top 20 most abundant genera for TCGA-BLCA as measured in this study, averaged over 129 WGS primary tumor and 27 solid tissue normal samples.

| Genus | Average read count per sample, this study | Average read count per sample, Poore *et al.* |
|---|---|---|
| *Enterococcus* | 447 | 9899 |
| *Veillonella* | 385 | 726 |
| *Staphylococcus* | 282 | 240880 |
| *Aerococcus* | 165 | 1756 |
| *Pseudomonas* | 133 | 44186 |
| *Peptoniphilus* | 95 | 737 |
| *Finegoldia* | 80 | 258 |
| *Stenotrophomonas* | 64 | 846 |
| *Anaerococcus* | 50 | 2451 |
| *Prevotella* | 49 | 1062 |
| *Streptococcus* | 36 | 560212 |
| *Cupriavidus* | 36 | 137 |

| | | |
|---|---|---|
| *Actinotignum* | 31 | 153 |
| *Citrobacter* | 25 | 695 |
| *Cutibacterium* | 24 | 0 |
| *Betapolyomavirus* | 22 | 0 |
| *Xanthomonas* | 22 | 15681 |
| *Erysipelatoclostridium* | 22 | 1.1 |
| *Campylobacter* | 18 | 2185 |
| *Eimeria* | 12 | 0 |

**Table S3**. Average read counts for the top 20 genera, ranked by weights assigned by the machine learning classifier using the "all putative contaminants removed" dataset and classifying BLCA primary tumor samples versus all other tumor types. Counts are averaged over 129 WGS primary tumor and 27 solid tissue normal samples.

| **Genus** | **Average read count, Poore *et al.*** | **Average read count, this study** |
|---|---|---|
| *Nitrospira* | 7.5 | 0 |
| *Elizabethkingia* | 460 | 0.1 |
| *Leptospira* | 3053 | 0 |
| *Campylobacter* | 2185 | 18 |
| *Histophilus* | 162 | 0 |
| *Capnocytophaga* | 56 | 0.2 |
| *Chelativorans* | 0 | 0 |
| *Sediminibacterium* | 2.6 | 0 |
| *Scardovia* | 0.6 | 0 |
| *Lysobacter* | 47 | 0.2 |
| *Stomatobaculum* | 0 | 0 |
| *Gallibacterium* | 20 | 0 |
| *Turicella* | 0.1 | 0 |
| Betaretrovirus | 0.2 | 0 |
| *Exiguobacterium* | 789 | 0 |
| *Wolbachia* | 291 | 0 |
| *Bacteroides* | 12013 | 7.7 |
| Alphapapillomavirus | 403 | 0.1 |
| *Hydrogenibacillus* | 17 | 0 |
| *Candidatus Stoquefichus* | 75 | 0 |

**Table S4.** Average read counts for the top 20 genera, ranked by weights assigned by the machine learning classifier using the "all putative contaminants removed" dataset and classifying BLCA primary tumor samples versus solid tissue normal samples. Counts are averaged over 129 WGS primary tumor and 27 solid tissue normal samples.

| **Genus** | **Average read count, Poore *et al.*** | **Average read count, this study** |
|---|---|---|

| | | |
|---|---|---|
| *Lachnoclostridium* | 18 | 0.2 |
| *Methylobacter* | 1.7 | 0 |
| *Marinitoga* | 157 | 0 |
| *Vibrio* | 34216 | 0.2 |
| *Crocinitomix* | 1.3 | 0 |
| *Flammeovirga* | 2.3 | 0 |
| *Desulfobacter* | 0.4 | 0 |
| *Paeniclostridium* | 148 | 0.1 |
| *Aliihoeflea* | 0 | 0 |
| *Cellulomonas* | 73 | 0.1 |
| *Candidatus Nitrosopelagicus* | 0.2 | 0 |
| Tymovirus | 10 | 0 |
| Betapartitivirus | 72 | 0 |
| *Microvirga* | 77 | 0 |
| *Salsuginibacillus* | 0.6 | 0 |
| Nepovirus | 5.7 | 0 |
| *Aeromonas* | 435 | 1.9 |
| *Klebsiella* | 21069 | 2.1 |
| *Pusillimonas* | 0.5 | 0 |
| *Candidatus Evansia* | 1.7 | 0 |

**Table S5**. Average number of reads in the top 20 most abundant genera for TCGA-HNSC as computed in the Poore *et al.* study, averaged over 334 WGS samples.

| **Genus** | **Average read count, Poore *et al.*** | **Average read count, this study** |
|---|---|---|
| *Streptococcus* | 1335308 | 2041 |
| *Mycobacterium* | 1001984 | 22.9 |
| *Staphylococcus* | 670494 | 121 |
| *Pseudomonas* | 273551 | 5685 |
| *Escherichia* | 212531 | 64.4 |
| *Mesorhizobium* | 158289 | 115 |
| *Waddlia* | 144680 | 0.0 |
| *Bacillus* | 137082 | 11.6 |
| *Neisseria* | 128526 | 1651 |
| *Pseudoalteromonas* | 116176 | 1.0 |
| *Streptomyces* | 96294 | 16.8 |
| *Vibrio* | 93027 | 2.8 |
| *Bordetella* | 84532 | 5.3 |
| *Shigella* | 70859 | 1.6 |
| *Piscirickettsia* | 66821 | 0.0 |
| *Treponema* | 63757 | 4774 |
| *Fusobacterium* | 57873 | 10003 |

| | | |
|---|---|---|
| *Salmonella* | 53463 | 1.2 |
| *Klebsiella* | 51586 | 34.5 |
| *Microbacterium* | 50001 | 10.1 |

**Table S6**. Average number of reads in the top 20 most abundant genera for TCGA-HNSC as measured in this study, averaged over 334 WGS samples.

| **Genus** | **Average read count, this study** | **Average read count, Poore *et al.*** |
|---|---|---|
| *Fusobacterium* | 10003 | 57873 |
| *Capnocytophaga* | 5981 | 13925 |
| *Prevotella* | 5706 | 47835 |
| *Pseudomonas* | 5685 | 273551 |
| *Treponema* | 4774 | 63757 |
| *Campylobacter* | 2102 | 26697 |
| *Streptococcus* | 2041 | 1335308 |
| *Neisseria* | 1651 | 128526 |
| *Veillonella* | 1328 | 3826 |
| *Haemophilus* | 1270 | 6870 |
| *Sphingomonas* | 1094 | 12979 |
| *Leptotrichia* | 721 | 1893 |
| *Stenotrophomonas* | 590 | 2646 |
| *Parvimonas* | 546 | 4062 |
| *Tannerella* | 428 | 2887 |
| *Porphyromonas* | 410 | 17431 |
| *Selenomonas* | 349 | 1111 |
| *Rothia* | 331 | 1027 |
| *Bifidobacterium* | 323 | 1522 |
| *Bradyrhizobium* | 301 | 2852 |

**Table S7**. Average read counts for the top 20 genera, ranked by weights assigned by the machine learning classifier using the "all putative contaminants removed" dataset and classifying HNSC primary tumor samples versus all other tumor types. Counts are averaged over 170 WGS primary tumor, 140 blood derived normal, and 24 solid tissue normal samples.

| **Genus** | **Average read count, Poore *et al.*** | **Average read count, this study** |
|---|---|---|

| | | |
|---|---|---|
| *Microvirga* | 209 | 1.8 |
| *Gemmata* | 13385 | 0.4 |
| *Desulfomicrobium* | 1.7 | 0.8 |
| *Exiguobacterium* | 1331 | 0.3 |
| *Nitrosopelagicus* | 0.4 | 0.0 |
| Alphapapillomavirus | 8577 | 0.4 |
| *Phenylobacterium* | 21 | 4.0 |
| *Klebsiella* | 51586 | 35 |
| *Plantibacter* | 0.3 | 0.3 |
| *Terracoccus* | 5.0 | 0.0 |
| *Marichromatium* | 0.8 | 0.3 |
| Betapartitivirus | 159 | 0.0 |
| *Gottschalkia* | 2.2 | 0.4 |
| *Acidithiobacillus* | 142 | 0.4 |
| *Desulfococcus* | 24868 | 0.2 |
| *Epilithonimonas* | 5.8 | 0.0 |
| *Luteibacter* | 1036 | 0.5 |
| *Spiroplasma* | 356 | 0.7 |
| *Mannheimia* | 429 | 2.2 |
| *Roseivirga* | 4.2 | 0.1 |

**Tables S8, S9, and S10** (separate files): These tables contains all read counts, reported at the genus level, from the bladder cancer (BLCA), head and neck cancer (HNSC), and breast cancer (BRCA) samples from TCGA. The tables include read counts for bacteria, archaea, and viruses. All samples were classified against a KrakenUniq database as described in Methods. **Table S8** (156 x 1063) has read counts for 156 BLCA samples, including 129 primary tumor and 27 solid tissue normal samples, filtered to remove human reads as described in Methods.
Overall, 1,063 genera were identified (i.e., contained at least one non-zero count) in the 156 samples. **Table S9** (334 x 1573) contains read counts for 334 HNSC WGS, including 170 primary tumor samples, 140 blood derived normal samples, and 24 solid tissue normal samples, filtered to remove human reads as described in Methods. Overall, 1,573 genera were identified (i.e., contained at least one non-zero count) across the 334 samples. **Table S10** (238 x 1,200) contains read counts for 238 BRCA WGS samples, including 114 primary tumor samples, 106 blood derived normal samples, 16 solid tissue normal samples, and 2 metastatic samples, filtered to removed human reads as described in Methods. Overall, 1,200 genera were identified (i.e., contained at least one non-zero count) across the 238 samples.

**Supplementary Data File 1**. List of all species contained in the Kraken database used in this study.
**Supplementary Data File 2**. List of all genera contained in the Kraken database used in this study.

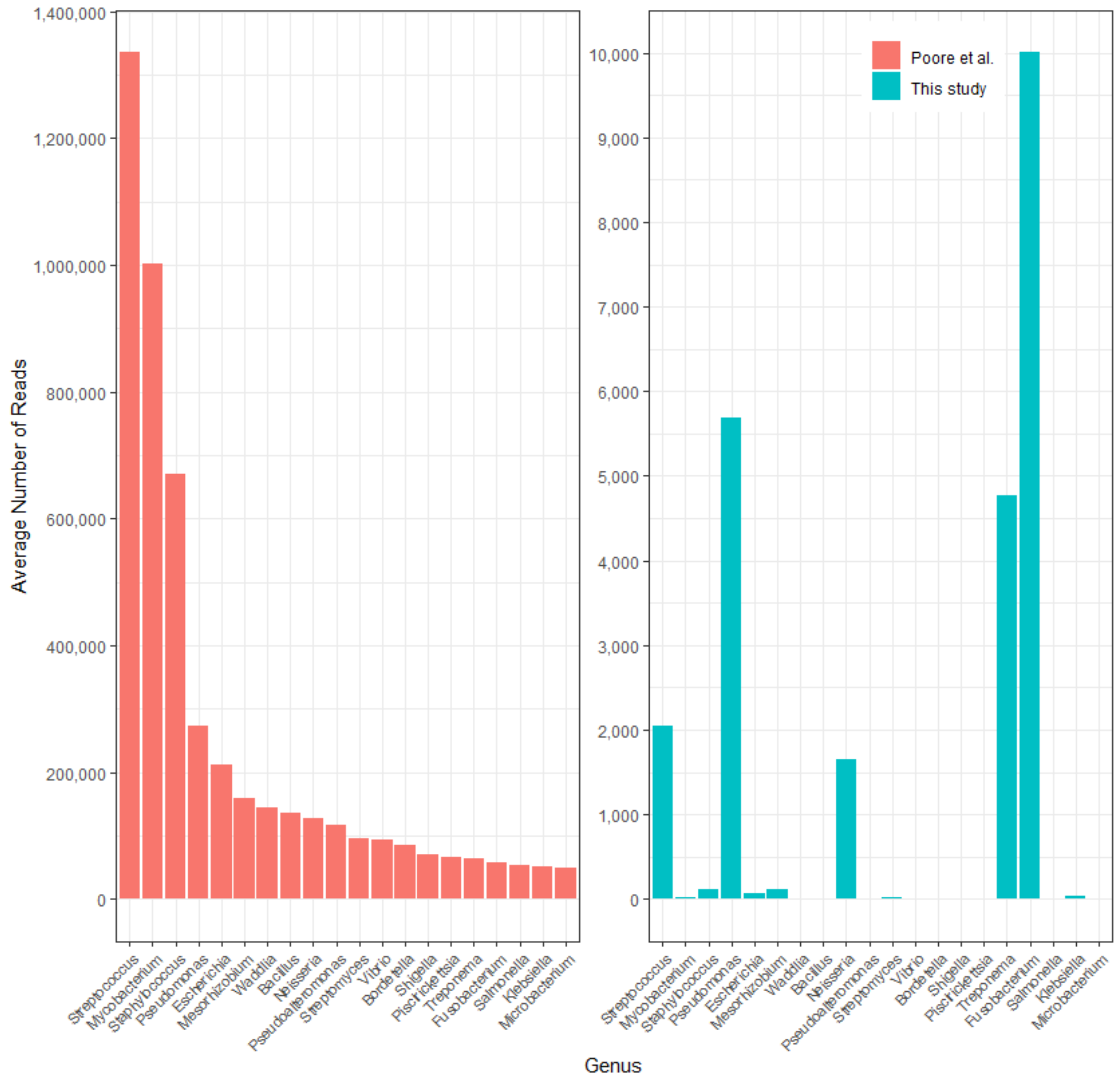


**Figure S1**. Average number of reads per sample in head and neck cancer (HNSC) in the top 20 most-abundant genera reported in Poore *et al*. (left), averaged over 170 WGS primary tumor, 140 blood derived normal, and 24 solid tissue normal samples. On the right are the counts for the same genera, in the same order, as computed in our re-analysis. Note that the y-axis scales are different by a factor of ~150. The x-axis shows genus names.

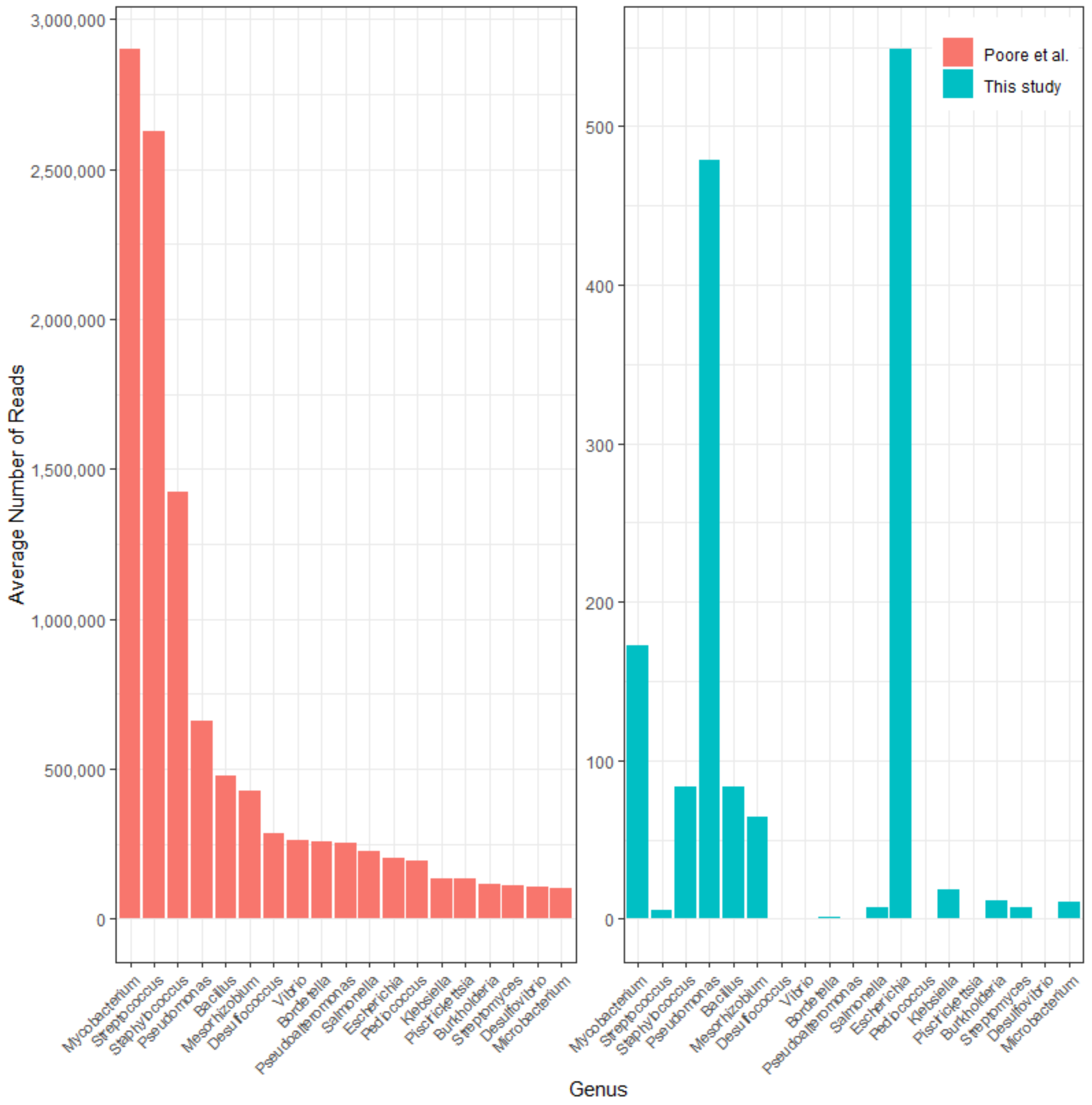


**Figure S2**. Average number of reads per sample in breast invasive carcinoma (BRCA) in the top 20 most-abundant genera reported in Poore *et al*. (left), averaged over 238 breast cancer WGS samples. On the right are the counts for the same genera, in the same order, as computed in our re-analysis. Note that the y-axis scales are different by a factor of ~5000. The x-axis shows genus names.
